# The evolution of open habitats in North America revealed by deep learning models

**DOI:** 10.1101/2021.09.03.458822

**Authors:** Tobias Andermann, Caroline Strömberg, Alexandre Antonelli, Daniele Silvestro

## Abstract

Open vegetation today constitutes one of the most extensive biomes on earth, including temperate grasslands and tropical savannas. Yet these biomes originated relatively recently in earth history, likely replacing forested habitats as recently as the second half of the Cenozoic, although the timing of their origination and the dynamics of their expansion remain uncertain. Here, we present a new hypothesis of paleovegetation change in North America, showing that open habitats originated between 25 and 20 Ma in the center of the continent, and expanded rapidly starting 8 Ma to eventually become the most prominent vegetation type today. To obtain space-time predictions of paleovegetation, we developed a new Bayesian deep learning model that utilizes available information from fossil evidence, geologic models, and paleoclimate proxies. We compiled a large dataset of paleovegetation reconstructions from the peer-reviewed literature, which we used in combination with current vegetation data to train the model. The model learns to predict vegetation based on the learned associations between the vegetation at a given site and multiple biotic and abiotic predictors: fossil mammal occurrences, plant macrofossils, estimates of temperature and precipitation, latitude, and the effects of spatial and temporal autocorrelation. Our results provide a new, spatially detailed reconstruction of habitat evolution in North America and our deep learning model paves the way for a new quantitative approach to estimating paleovegetation changes.

## Introduction

The different types of vegetation and their spatial distribution form the biotic landscape on which other species, including all terrestrial animals, interact and evolve. From reconstructions of past vegetation (paleovegetation) at different sites, we know that vegetation evolves and changes dynamically through time as it responds to environmental alternations, such as changes in climate ^1,2^, interaction with faunal communities ^3^, and large biological events such as mass extinctions ^4^.

Several major vegetation changes are documented in the fossil record, including the shift from ecosystems dominated by free-sporing plants to seed plants ^5^ and the radiation and ecological expansion of angiosperms ^4,6,7^, which today dominate most terrestrial biomes. The most important vegetation change in the Cenozoic is arguably the origination and expansion of open, grass-dominated habitats ^8^ at the expense of closed forest ecosystems ^9^. Open grasslands today represent the most extensive terrestrial biome on Earth, covering over 40% of the Earth’s land surface ^10^. The oldest confirmed presence of open-habitat grasses in North America dates back to the Late Eocene, yet the fossil record indicates that these were rare elements of the vegetation and at the time did not constitute sizable open ecosystems ^9,11^. Based on the currently available paleobotanical evidence it is likely that open grass-dominated habitats first appeared as a novel biome comparably recently in the Late Oligocene to Early Miocene ^8,12^, yet the dynamics of their expansion are still poorly understood and much debated (e.g., see ^9^).

Previous studies have produced paleovegetation reconstructions for individual sites based on the evaluation of i) plant macrofossil assemblages (i.e., fossilized leaves, seeds, wood, or other plant organs); ii) fossilized pollen; or iii) phytoliths —microscopic silica bodies produced in plant cells with a high fossilization potential and unique shapes, which can be attributed to specific vegetation components ^13^. While such reconstructions can provide an accurate record of the vegetation at a given site, extrapolating these reconstructions to larger geographic scales and through time is hampered by the sparsity of fossil sites and the incompleteness of the record. Such extrapolations can be done based on expert opinion under consideration of paleoclimatic models and other information ^14–16^, at the cost of reduced reproducibility and limited scalability.

Most modeling studies that are aiming to infer past or future vegetation have been based on climate models and predefined tolerance limits for certain biome types ^17,18^. However, additional types of data hold great potential for such modeling approaches, such as the associations between the faunal fossil record with the surrounding vegetation. For example, the relationship between grassland biomes and large grazing mammals, often identifiable by their hypsodont teeth, has long been used to infer presence of grasslands ^19,20^ (but see Dunn et al. ^21^). Such information about plant-mammal interactions is commonly used to manually infer the paleovegetation at individual fossil sites ^9^, and sometimes mammal fossil assemblages are used to validate and correct vegetation predictions made from climate-based models ^18^.

Mammal fossils are a useful data type because of their relatively rich record. Further, mammals are one of the paleontologically best studied groups with a relatively well-resolved fossil taxonomy, often allowing for precise identifications of fossil mammals down to genus or even species level. Many of these mammal fossil data are publicly available through large online databases (e.g., Alroy et al. ^22^). Yet, to our knowledge, no computational models exist that explicitly utilize this information to predict vegetation. This is partly because for most taxa, the habitat associations are difficult to establish with confidence, particularly so for extinct taxa, and ambiguous for many mammal taxa that are not restricted to a single vegetation type.

To seize the opportunities outlined above and improve the reconstruction of past environments, we present here a Bayesian deep neural network (BNN) model that utilizes the available information on mammal fossils, plant macrofossils, modeled paleoclimate data, as well as spatial and temporal associations to predict vegetation. Importantly, our model does not require any prior assumptions on temperature tolerance limits or ecological interactions. Instead, it learns how these biotic and abiotic features can be mapped to a vegetation type within a supervised learning framework. This property provides great flexibility, as any available biotic or abiotic predictor can be added to the model, while the model decides based on the data whether this predictor is deemed useful for the vegetation inference. Once trained, the model estimates the most likely vegetation for any given point in time and space, and the uncertainty associated with the prediction. We demonstrate the utility of our methodology by modeling past vegetation changes in North America throughout the last 30 million years (Ma), focusing on the expansion of open, grass-dominated habitats.

## Results

### Model description

We implement a BNN model to predict vegetation through space and time (Fig.1, see Methods for a more detailed description). We focus on two vegetation types “open vegetation” (open grasslands, savannas and steppes, desert vegetation, and tundra) and “closed vegetation” (forests); additional categories could be implemented for other systems if sufficient data are available for training. As features for this classification task we use biotic data, consisting of fossil occurrences of 100 selected mammal and plant taxa (see Methods), supplemented by current occurrences of these taxa. Further, we use several abiotic features including proxies for climatic data (mean annual temperature and precipitation) through space and time ^23^, paleocoordinates ^24^, and mean global temperature from oxygen isotope data ^25^.

**Figure 1:**
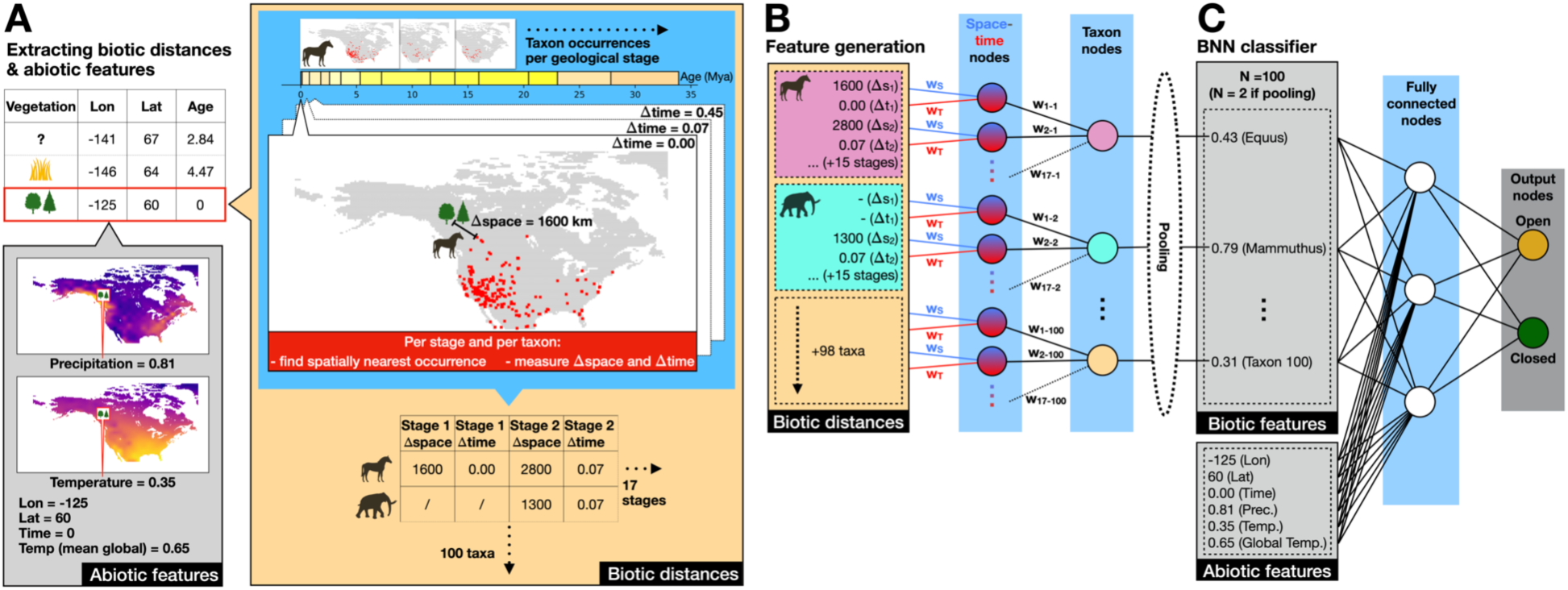
The process of feature generation and the BNN model architecture applied in this study. (A) For a given point defined by its longitude (Lon), latitude (Lat), and age, here highlighted in red, we extract the spatial distance to the closest occurrence of each taxon. This is repeated for each geological stage (n=17), while also extracting the temporal distance between the given point and the mid age of each geological stage. (B) These spatial and temporal distances are the input of the first two hidden layers in our BNN for feature generation. The BNN estimates weights (red and blue lines) to reduce the multitude of spatial and temporal distance measurements into one single “proximity” value for each taxon (taxon nodes). (C) These taxon features (biotic features) are then used in combination with abiotic features, reflecting climatic, geographic, and temporal variables, as input into the fully connected BNN classifier. During training the weights (black lines connecting nodes) of the feature generation layers and of the BNN classifier are estimated through MCMC sampling to optimally map the input data to the correct output vegetation label (“open” or “closed”). Once trained, a posterior sample of the weights is stored for each model and is used to make vegetation predictions for points with unknown vegetation.

Our deep neural network consists of two pre-processing layers, where distances derived from the raw mammal and plant fossil occurrence data (Fig. 1A) are transformed into taxon-specific features (Fig. 1B, see “Feature generation” in Methods section for a more detailed explanation). These features comprise the input data for the fully connected classification layers of our BNN model, which quantify the probability of each vegetation type (Fig. 1C). Through this setup the model is trained to infer the vegetation type based on the measured distances to nearby taxon occurrences, in combination with the additional climatic and geographic features.

### Predicting past vegetation

To train our BNN vegetation classifier, we compiled a total of 331 paleovegetation reconstructions based on phytolith and pollen assemblages, paleosol data and macrofossils from the peer-reviewed literature (see Methods), ranging in age from the beginning of the Oligocene (33.9 Mya) to the present (Supplementary Fig. S2). To further increase the number of training data, we supplemented the paleovegetation data with data about current vegetation. Since current vegetation patterns are heavily influenced by human activity, we retrieved the SYNMAP Global Potential Vegetation data, representing the potential vegetation without human land alterations ^26^. To find the model configuration that produced the best paleovegetation prediction accuracy, we tested a range of different model architectures, as well as different combinations of input data (Table 1). We applied a five-fold cross validation approach to the paleovegetation data when training each model; in this approach each model is trained five times on a different 80% of the input data, while using the remaining 20% as a test set. This allows to determine the overall prediction accuracy of the model by averaging the number of correctly predicted test set labels across all 5 test sets, comprising all available data. We calculated the prediction accuracy separately for paleovegetation and current vegetation, as well as a combined weighted mean of the two (Table 1, Supplementary Fig. S1, see Methods for more information).

**Table 1.** Prediction accuracy of tested model configurations. The overall accuracy of each model constitutes the weighted mean between the paleovegetation accuracy (weighing-factor 10) and the current vegetation accuracy (weighing-factor 1). This weighing was applied because the paleovegetation data were distributed across 10 geological stages, while the current accuracy only represents one single stage. The selected posterior probability (PP) threshold was chosen to reach a minimum prediction accuracy of 90% on the weighted test set. For some models this accuracy aim could not be achieved; in these cases, the posterior threshold was set to 1, leading to all vegetation predictions to be labeled as ‘unknown’, when applying this threshold.

| Model ID | Architecture | Train instances | Features | Pooling | Accuracy | Accuracy (paleo) | Accuracy (present) | Selected PP threshold | Fraction of predictions above PP threshold |
| --- | --- | --- | --- | --- | --- | --- | --- | --- | --- |
| 1 | 2 layers, 32-8 nodes | 281 current, 281 paleo | All | None | 0.888 | 0.891 | 0.863 | 0.620 | 0.970 |
| 2 | 2 layers, 32-8 nodes | 281 current, 281 paleo | Only biotic | None | **0.892** | **0.894** | 0.872 | 0.570 | 0.970 |
| 3 | 2 layers, 32-8 nodes | 281 current, 281 paleo | Only abiotic | None | 0.870 | 0.870 | 0.874 | 0.630 | 0.937 |
| 4 | 2 layers, 32-8 nodes | 281 current, 281 paleo | All | Sum | 0.883 | 0.885 | 0.858 | 0.630 | 0.934 |
| 5 | 2 layers, 32-8 nodes | 281 current, 281 paleo | Only biotic | Sum | 0.883 | 0.885 | 0.862 | 0.570 | 0.940 |
| 6 | 1 layer, 8 nodes | 281 current, 281 paleo | All | Sum | 0.879 | 0.879 | 0.879 | 0.590 | 0.943 |
| 7 | 1 layer, 8 nodes | 281 current, 281 paleo | Only biotic | Sum | 0.881 | 0.882 | 0.872 | 0.570 | 0.940 |
| 8 | 1 layer, 8 nodes | 281 current, 281 paleo | Only abiotic | None | 0.817 | 0.825 | 0.735 | 0.850 | 0.239 |
| 9 | 2 layers, 32-8 nodes | 281 current, 281 paleo | All | Max | 0.886 | 0.888 | 0.866 | 0.640 | 0.921 |
| 10 | 2 layers, 32-8 nodes | 281 current, 0 paleo | All | None | 0.516 | 0.480 | 0.881 | 1.000 | 0.000 |
| 11 | 2 layers, 32-8 nodes | 1405 current, 0 paleo | All | None | 0.683 | 0.660 | **0.916** | 1.000 | 0.000 |
| 12 | 2 layers, 32-8 nodes | 0 current, 281 paleo | All | None | 0.855 | 0.876 | 0.647 | 0.640 | 0.909 |
| 13 | 2 layers, 32-8 nodes | 1405 current, 281 paleo | All | None | 0.841 | 0.834 | 0.906 | 0.680 | 0.846 |

The best model (#2, Table 1) reached a prediction accuracy of 89.2% (89.4% paleo, 87.2% current). The model included only the biotic features (taxon distances) and its architecture consisted of two layers containing 32 and 8 nodes, and no feature pooling (see Methods for more details). The prediction accuracy can be further improved by applying a posterior probability (PP) threshold to the class predictions, only making vegetation inferences for predictions that exceed this threshold (see Supplementary Results for more details). The higher the PP threshold is set, the higher test accuracies can be reached, at the cost of an increasing number of test instances being predicted as “unknown”, as they fall below the threshold (Fig. 2). Here, we determine a PP threshold for each of our trained models to ensure a minimum test accuracy of 90%. For the best model, a PP threshold of 0.57 was sufficient to reach an expected target accuracy of >90%, while still making vegetation inferences for 97% of the test set (Table 1).

**Figure 2.**
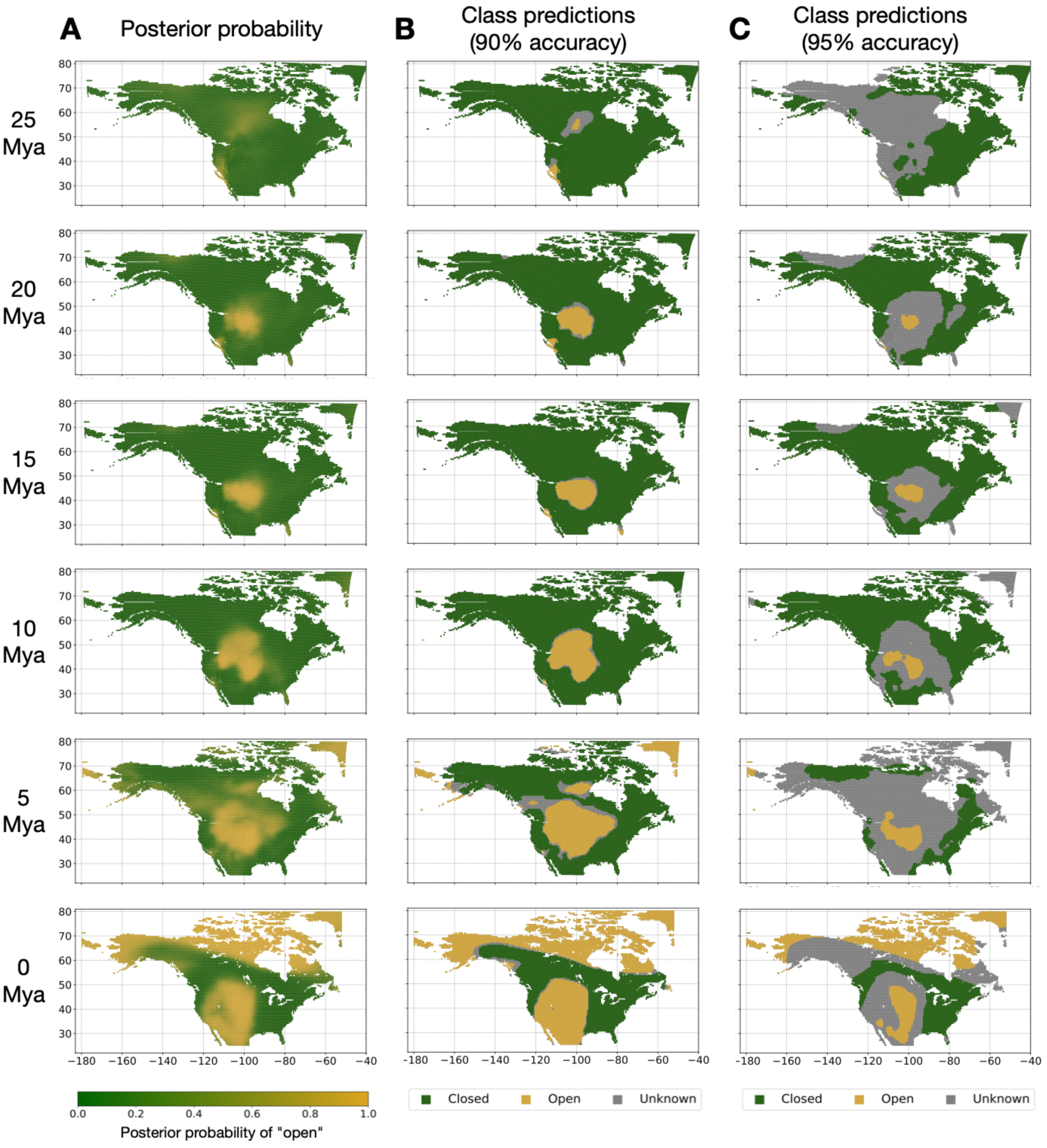
Vegetation predictions for North America throughout the last 30 Mya. The predictions are based on the best model resulting from our model evaluation and sensitivity tests (#2, Table 1). Panel A shows the posterior probability (PP) estimates for open habitat, where a PP of > 0.95 (yellow) indicates strong evidence for open habitat, whereas a PP of < 0.05 (green) indicates strong evidence for closed habitat. Panels B and C show categorical vegetation class predictions for our vegetation classes “open” (yellow) and “closed” (green). The class predictions are based on a PP threshold ensuring 90% prediction accuracy (B), and 95% prediction accuracy (C), respectively. The higher the applied PP threshold, the more sites will be classified as “unknown” (grey).

Using the best of our trained models, we produced vegetation estimates for North America throughout the last 30 Mya in 1 Mya increments. To further improve the model for predicting past vegetation, we retrained it using all available paleovegetation points, including those 20% that were previously used for model evaluation. The final model was trained on all 331 paleovegetation points and 281 current vegetation points. To produce the data for the prediction task, we calculated the cell center coordinates of all land cells in a 0.5° x 0.5° grid across the majority of the North American continent, which we defined by a cropping window with corner points P_1_ (Lon = -180, Lat = 25) and P_2_ (Lon = -52, Lat = 80). We accounted for tectonic movements, transforming the grid-cell center coordinates into their equivalent paleocoordinates, using the “PALEOMAP” model of the mapast R-package ^24^. From the grid-cell center coordinates, we extracted the taxon occurrence distances for feature generation, as well as all additional abiotic features, in the same manner as for the training and test data (Fig. 1).

Our inferred paleovegetation maps suggest that small pockets of open habitats may have already existed in North America around the end of the Oligocene epoch (25 Mya, Fig. 2), but these only constituted a small fraction of the total vegetation (<10%, Fig. 3). Open habitats stayed constant (<20% of total vegetation cover) or expanded slowly throughout most of the predicted time frame. Only toward the late Miocene at about 8 Mya did open habitats begin to increase rapidly, at present making up >60% of the North American natural vegetation (estimated in the absence of anthropogenic impact; Fig. 3). These patterns of open habitat expansion are predicted consistently across different models tested (Fig. 3).

**Figure 3.**
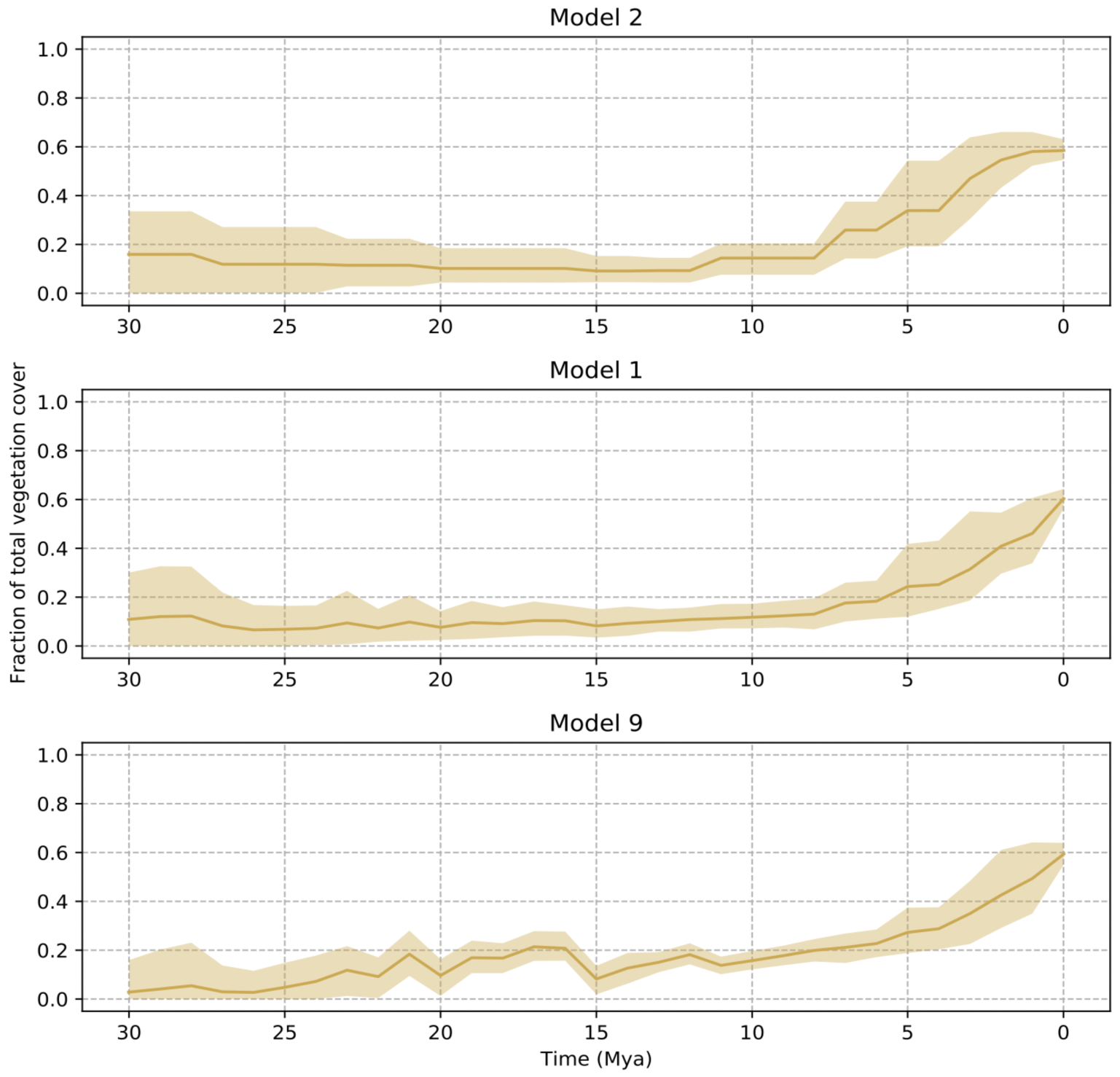
Proportion of predicted open habitat through time, across all terrestrial cells, for the three best models (#2, #1, and #9) resulting from our model testing (Table 1). Fractions are calculated as the proportion of all terrestrial cells. The solid line shows the mean estimates of open habitat fraction across all posterior samples, while the shaded area shows the 95% HPD interval. All displayed models show overall similar patterns (ignoring minor fluctuations) of open habitat expansion in North America, with open habitats existing at low frequency (<20%) until their continuous expansion starting approximately 8 Mya.

To determine to what degree the mammal taxa (genera) used in this study are associated with either open or closed vegetation, we evaluated for each taxon the fraction of occurrences that fall within each vegetation type, based on the vegetation predictions made by our best model. The open habitat specialist mammal genera identified by our predictions are *Cynomys, Onychomys, Ochotona, Perognathus, Zapus*, and *Brachyerix*, which all have more than 75% of their occurrences through time inside of grid cells with inferred open vegetation (Supplementary Table 1). On the other hand, we identified *Dasypus, Lontra, Panthera, Eumops, Sylvilagus, Erethizon*, and *Gompotherium* as the most specialized closed vegetation mammal genera (>75% closed vegetation occurrences). This demonstrates one utility of our model, as it allows us to estimate mammal-vegetation associations from the predictions made by the trained model, rather than having to define these a-priori.

### Sensitivity tests

We tested whether the prediction accuracy of our models improves by supplementing the paleovegetation data with current vegetation data when training the model. We found that in fact the prediction accuracy for both, paleovegetation and for current vegetation, increases when adding small numbers (n=281, Supplementary Fig. S3) of current vegetation datapoints during model training (see model #1 vs. #12, Table 1). The prediction accuracy for current vegetation further improves when adding increasing numbers of current vegetation instances (n=1,405, Supplementary Fig. S3), yet at the same time the prediction accuracy for paleovegetation starts to decline (model #1 vs. #13, Table 1). This indicates that our model utilizes some of the information that is gained from adding current vegetation information during training, but that it overfits toward the present when the number of current vegetation instances exceeds the number of paleovegetation instances by a multitude (factor 5 or higher).

Further, we tested whether the addition of biotic features (taxon distances) improved the prediction accuracy, compared to models using only abiotic features, such as those used in previous studies based primarily on climate ^17,18^. In fact, we find model #2, which only utilizes biotic features, to be the overall best model, outperforming the model only based on abiotic features (model #3), and even outperforming models based on both biotic and abiotic features by a small margin (model #1). Further analyses of feature importance show that indeed several of the biotic mammal features (taxa) stand out as the features with the highest impact on the prediction accuracy of the model (in particular the genera *Castor* and *Mammut*) and that abiotic features provide only a limited contribution toward the prediction accuracy (Fig. 4). This finding suggests that mammal and plant fossil occurrence data capture most of the relevant information to predict paleovegetation changes.

**Figure 4.**
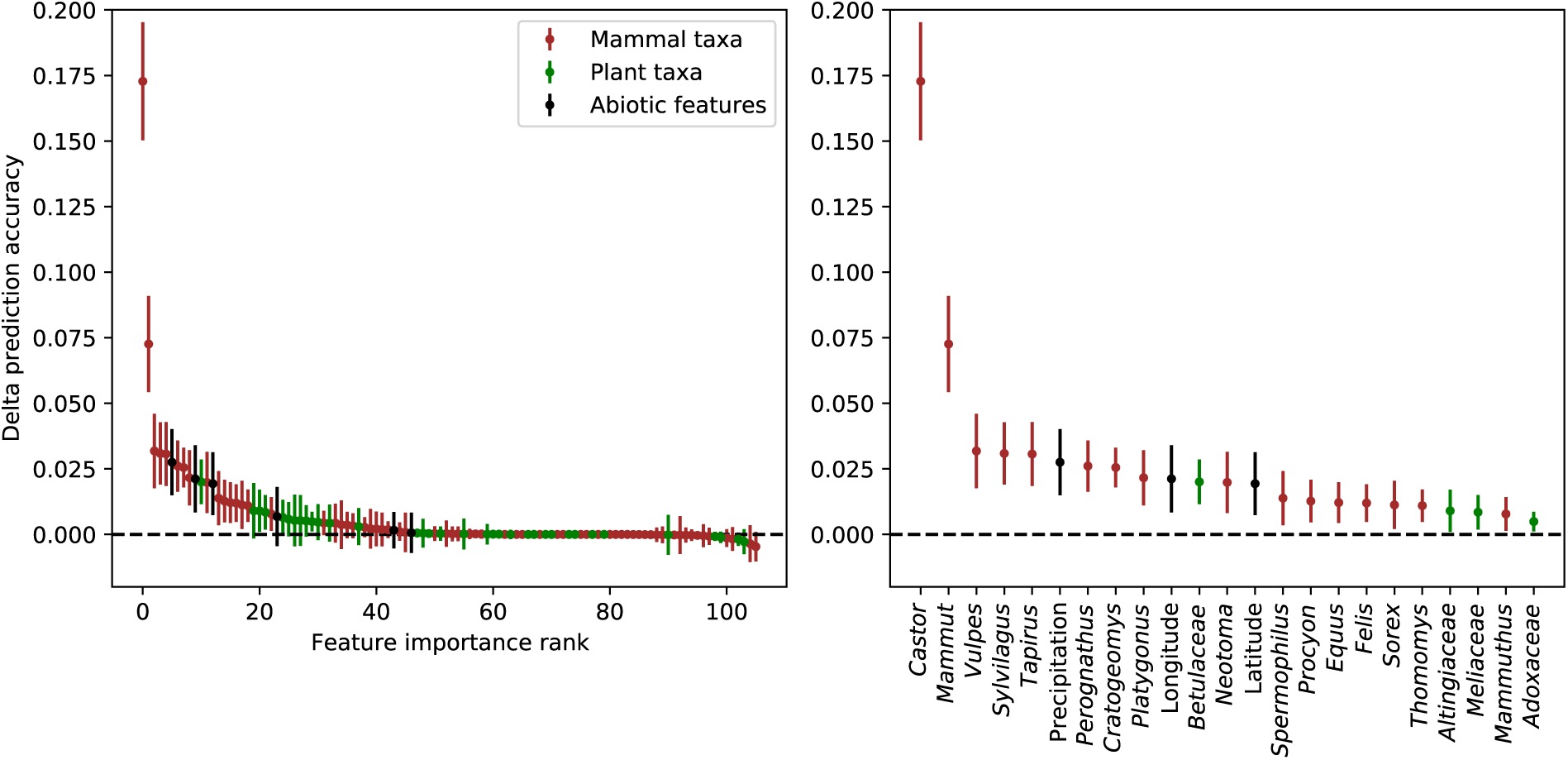
Impact of individual features on model prediction accuracy. The displayed delta-accuracy values for each feature reflect by how much the prediction accuracy of the trained model drops if the information content of a given feature was removed. This is accomplished by randomly shuffling the values of a given feature across all test set instances, and then subtracting the resulting prediction accuracy of this modified test set from that of the unmodified test set (permutation feature importance). Points show the mean delta-accuracy of each feature across 100 randomly selected posterior BNN weight samples, while the error bars show the standard deviation. The first panel shows the delta-accuracy estimates for all 106 features, while the second panel shows only those features with a consistently positive feature importance across all posterior samples, representing the most important features for the trained model.

Finally, we tested different approaches of feature reduction by pooling the numerous biotic features (n=100) into one single faunal and one single floral feature (n=2), before feeding them into the fully connected BNN layers for classification (Fig. 1). This approach of pooling greatly reduces the number of weights that need to be estimated, leading to faster training and better convergence of the MCMC used to sample the BNN weights. While this did not lead to improvements in the prediction accuracy, these models generally performed equally well compared to the more complex models, allowing us to reach similar prediction accuracies with a substantially decreased number of parameters (model #6 vs. #1, Table 1).

## Discussion

Here we present a probabilistic reconstruction of paleovegetation and its evolution for North America, based on a deep learning model. This model paves the way for a universally applicable, full evidence approach, which utilizes raw geographic and temporal distances to taxon occurrences, in conjunction with abiotic data such as climate and spatiotemporal coordinates. The spatial and temporal distances that are required as input can be easily calculated for any given point in space and time, independent of its vicinity to the nearest fossil record of a given taxon, which makes our model applicable to a wide range of geographic and temporal contexts.

Our trained models predict the presence of comparably small pockets of open habitat at the beginning of the Oligocene-Miocene transition, existing at low frequencies until the end of the Miocene, when open habitats started increasing at a continuous pace to finally reach their present extent (Figs. 3 and 4). These model predictions support a scenario of comparably late spread of open grasslands across vast areas in North America. It can be compared to the two-stage model for grassland expansion in the Great Plains based on primarily phytolith data, whereby woodlands and mosaics of the Early-Middle Miocene were replaced by open grasslands in the Late Miocene ^9,27^—the major novelty here being that our study adds a high-resolution geographic and temporal dimension.

Other proposed scenarios, based on the fossil pollen record and macrofossils, place the expansion of open habitats in North America much earlier, in the Middle Eocene, although these early open habitats were likely not grassland dominated ^28^. Similarly, paleovegetation reconstructions based on paleosols suggest the presence of open habitat grasslands already in the Late Eocene ^29^. While each of these previous reconstructions are based on a different type of paleovegetation data, our predictions are based on a general representation of the overall paleobotanical evidence, as they were trained on a mix of phytolith, pollen, macrofossil, and paleosol data (Supplementary Data S1), as well as current vegetation information. The results of our cross-validation approach reflect how well each model can learn from and re-predict the entirety of these input data; the fact that our best model reaches a prediction accuracy of >89% (Table 1), demonstrates that these heterogeneous data types can be meaningfully combined in a mixed evidence model.

In this study we restricted the model to only two broad vegetation classes, “open” and “closed”, due to limited availability of paleovegetation data points for training. More concerted efforts are needed to produce and compile larger and more spatially complete paleovegetation datasets based on pollen, phytoliths, or macrofossil assemblages to ensure sufficient training data for more detailed inferences of paleovegetation. This would allow predicting more nuanced vegetation types, for example distinguishing between taiga, temperate and tropical forest, as well as between tundra, temperate grasslands, and tropical steppes.

Given the coarse temporal resolution due to the binning of our data into geological stages, the models trained in this study are not appropriate to infer the influence of short-term climatic fluctuations, such as recent glaciations and the corresponding vegetational changes ^30^. However, our deep learning method could be used with high-resolution temperature and precipitation data in combination with the detailed Quaternary fossil record, to predict recent vegetation changes at spatiotemporally finer scales. For example, such models could be applied to predict vegetational changes linked to the Quaternary glacial cycles and to the human expansion and megafauna extinction (e.g., ^31,32^). Further, variations of the model developed here could also be applied to forecast future vegetational changes based on predictions of climate and other environmental factors.

## Methods

Here we estimate vegetation patterns through time and space for North America. For this purpose, we develop and train a (BNN) model to learn the associations between sites with current or past vegetation information and nearby mammal and plant fossil occurrences, as well as associations with climatic and spatiotemporal factors. We then apply the trained BNN to estimate vegetation labels for spatial grids throughout the last 30 Ma, to estimate vegetational changes in North America through time and trace the evolution of grass-dominated open habitats.

### Data

#### Spatial and temporal range

In this study we focused on a geographic area that is defined by a cropping window with the corner points P_1_ (Lon = -180, Lat = 25) and P_2_ (Lon = -52, Lat = 80), covering the majority of the North American continent (Figure 3). The temporal focus of this study encompasses the last 30 Ma, which is the time span containing the majority of our available sites with paleovegetation information (Figure S1). From the following data sources, we only selected those data points that fall within this spatiotemporal range.

Our approach described below required discretizing the input data of past vegetation labels and fossil occurrences into time-bins. For this we chose the age boundaries of geological stages defined in the International Chronostratigraphic Chart, v2020/03 ^33^, since these stages are expected to represent meaningful temporal units for analyzing both faunal and floral patterns. A total of 17 geological stages fell within our selected time frame of the last 30 Ma. We discretized the ages of all data points (vegetation data and fossil occurrences) that fell within a given stage by setting them to the midpoint of the respective stage.

#### Paleovegetation data

We reviewed a large body of peer-reviewed literature containing paleovegetation reconstructions and compiled a database of 1,242 paleovegetation data points for North America (Supplementary Data S1). These data represent individual vegetation reconstructions based on fossil evidence (phytoliths, pollen, macrofossil assemblages). We condensed the vegetation interpretation of the compiled vegetation data, which in many cases described specific vegetation ecosystem components, into the broader labels “open” versus “closed” habitat. In several cases, we found multiple vegetation reconstructions for the same spatiotemporal site, for example when multiple sediment samples were taken from the same horizon of a given formation. We treated these spatiotemporal duplicates as a single data point, resulting in a total of 331 spatiotemporally unique paleovegetation data points, of which 180 were labeled as “closed” and 151 as “open” (Supplementary Data S1). In the few cases (n=3) where both “open” and “closed” vegetation interpretations were present for the same spatiotemporal point, we selected one at random.

#### Current vegetation data

To supplement the limited number of paleovegetation sites, we compiled data about the current vegetation within our study area. In order to obtain current vegetation patterns, we downloaded the SYNMAP Global Potential Vegetation data ^26^. As for the paleovegetation data, we collapsed the more detailed biome data into broader categories by coding biome IDs < 37 as “closed” and biome IDs ≥ 37 as “open”. The resolution of the resulting raster was 0.5° longitude x 0.5° latitude, which equates to a spatial resolution of approximately 50 × 50 km grid cells (at the equator). We extracted all current vegetation grid cells that fell within our defined cropping window (Figure 3), excluding all sea water cells as well as large continental lakes. This resulted in 11,048 terrestrial grid cells with current vegetation information. For these grid cells we extracted the coordinates of the cell-center as well as the corresponding vegetation label.

The compiled paleovegetation and current vegetation points constitute the pool of vegetation information from which we sampled subsets to train our model. From here on we refer to these data points as our training instances. We trained several BNN models, using different combinations of the paleovegetation points (n = 331) and current vegetation points (n = 11,048).

#### Fossil data

We downloaded all available mammal fossil data from major public databases, namely the Paleobiology Database ^22^, the NOW (New and Old Worlds) database of fossil mammals ^34^, the Neotoma database ^35^, and the Miocene Mammal Mapping Project ^36^. We merged all downloaded fossil occurrences into one shared database and removed all entries that were not identified to species level, as well as all spatiotemporal duplicates. In several cases the fossil data downloaded from the major databases contained minor spelling inconsistencies in the genus names and species epithets. To correct these misspellings, which can lead to an overestimation of the number of genera and species in the dataset, we used the algorithm implemented in the PyRate package ^37^, which automatically identifies common typos in scientific names. Finally, we removed all aquatic families from the dataset (dugongs, pinnipeds, and whales).

For each fossil occurrence we determined the mean age of the respective stratigraphic age interval. We reduced the taxonomic resolution of the mammal data to genus-level with the main purpose to reduce the number of taxa, while increasing the spatial and temporal extent of each taxon as well as to avoid taxonomic biases (such as over-splitting or lumping of species in different genera, depending on taxonomic authority). Both of these issues are expected to have a smaller impact on genus level compared to species level. To further reduce the number of taxa to only the most informative ones, we only kept genera that were present in more than half of the geological stages covered in this study, based on the first and last occurrence date of each genus in the fossil record (assumed presence in at least 9 of 17 stages). This resulted in 65 selected mammal genera (Supplementary Table S1). While the model can potentially handle any number of taxa, taxa with occurrences spanning multiple locations and time bins are expected to be most informative in our supervised learning approach.

As an addition to the mammal fossil data, we compiled a large dataset of plant macrofossils from the Cenozoic Angiosperm database ^38^. Due to taxonomic inconsistencies of fossil plants on species and genus level, we decided to reduce the taxonomic resolution of the plant fossil data to family level. Similar to the mammal fossil data, we took the mean age of the stratigraphic age interval of each fossil occurrence and only selected plant families that were present in North America during at least 9 of the 17 geological stages. This resulted in 35 selected plant families (Supplementary Table S1). The final fossil data, consisting of the selected mammal and plant taxa (n=100), amounted to a total of 7,501 selected fossil occurrences (6,759 mammal and 742 plant fossils, Supplementary Data S2).

#### Current occurrences

To complement the occurrence data extracted from the fossil record, we extracted current occurrences for all selected taxa from the Global Biodiversity Information Facility (GBIF) ^39^. For all mammal genera we downloaded the data through the R-package rgbif ^40^, only allowing human observations (as opposed to e.g. machine observations or fossil data) and restricting the search to North American occurrences (Canada, Mexico, or USA), using the following command:

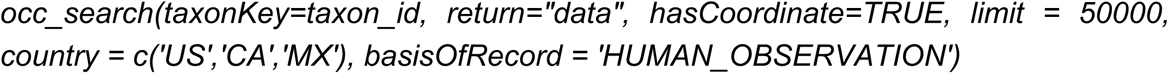

Due to the large data volumes for the selected plant families, which result in very long waiting times and occasional time-out errors when using the rgbif package, we instead downloaded the current occurrences of the selected plant families directly from the GBIF online interface (download DOI: https://doi.org/10.15468/dl.nxuyg8). This resulted in a total of 1,251,810 current occurrences for the selected extant mammal and plant taxa (103,813 mammal and 1,147,997 plant occurrences, Supplementary Data S2). Finally, all fossil and current occurrences of the selected taxa were merged into one data-frame and jointly treated as occurrence data, independently of the data origin as fossil or GBIF observation. For all further steps, we only selected those occurrences that fell within the cropping window defined as described above.

#### Climatic data

We downloaded global maps of modeled precipitation and temperature from Scotese et al. ^23^, with a spatial resolution of 1° longitude x 1° latitude. Because these data are only available in 5 Ma intervals, we linearly interpolated the values into 1 Ma year intervals to reach higher temporal resolution. Additionally, we downloaded estimates of mean global temperature that are based on oxygen isotope data ^25^.

### Feature generation

An essential element of applying neural networks is the process of feature generation, which describes the transformation of the raw data into numerical features that can be fed into the neural network. Each input data point, which is commonly referred to as an instance, consists of a list of associated feature values. In our case the training instances consist of specific points in space-time with available vegetation information, and the associated features contain the information about nearby occurrences of the selected taxa (biotic features), as well as other information about climate, geography, and time (abiotic features), in relation to the given point.

#### Biotic features

For a given instance (vegetation point), defined by its spatial and temporal coordinates, we extracted the geographic distance (Mercator distance) between this instance and the closest occurrence of each taxon, and we did so for each geological stage (Fig. 1). If a taxon was present in all stages, this resulted in 17 geographic distances extracted for this taxon, one for each stage. These spatial distances were calculated using the current coordinates (instead of the paleo-coordinates) of each point, assuming that the relative spatial distance between any two given points within North America is not affected (or negligibly so) by continental movements during the last 30 Ma, although their absolute coordinate values have changed through time.

Additionally, we extracted the temporal distances between the selected taxon-occurrences and the given vegetation point, by measuring the difference between the age of the training instance and the midpoint of the geological stage of a given taxon occurrence. This resulted in *N* pairs of geographic and temporal distances to each taxon, where *N* is the number of stages this taxon occurred in. We designed our BNN model to estimate parameters to summarize the spatial and temporal distances of the selected occurrences of each taxon into one taxon-specific feature value, representing a measure of general “proximity” of each taxon, which we explain in more detail below (Fig. 1).

#### Abiotic features

In addition to the biotic features, we extracted the temperature and precipitation associated with the space-time coordinates of a given instance. For this step we transformed the coordinates of each given vegetation label into the equivalent paleo-coordinates at the time of the record, using the “PALEOMAP” model of the *mapast* R-package ^24^. We extracted the modeled temperature and precipitation of these paleo-coordinates from our rasterized climate data as two separate features. As an additional climatic feature, we extracted the mean global temperature at the given time point. Finally, we added the absolute paleo-coordinates (longitude and latitude) as well as the given time of the vegetation point as three additional features.

Our neural network was trained on a total of 100 biotic features (one for each selected taxon), 3 climatic features, and 3 spatiotemporal features, resulting in a total of 106 features for each instance.

To avoid potential biases based on the absolute values of given features, we scaled all features to a range between 0 and 1. The rescaling was done jointly for all training and prediction instances, in order to avoid differences in rescaling-factors between features in the training instances and those in the prediction instances.

### Selecting training and test data

For training our neural network we had a total of 11,379 points with vegetation information available, consisting of 331 paleovegetation points and 11,048 current vegetation points. To test whether the larger number of current vegetation instances might bias our past vegetation reconstructions, we explored different combinations of paleovegetation and current vegetation instances during training of the model (Table 1).

To evaluate the prediction accuracy of our trained models, we performed a five-fold cross validation, training each of the five cross validation models on 80% of the available paleovegetation instances, while sparing the remaining 20% as a test set. The current vegetation instances that were used for training some of our tested models remained the same across all cross-validation folds. We then determined the paleovegetation prediction accuracy of the model as the average test set prediction accuracy across all 5 cross validation folds. Additionally, we determined the prediction accuracy for all remaining current labels that were not used for training. The final prediction accuracy of each model was then determined as the weighted mean between the paleovegetation prediction accuracy and the current vegetation prediction accuracy of the model, weighing the paleovegetation component ten times higher, as it represents the accuracy across ten geological stages that are covered by our paleovegetation data (Supplementary Fig. S2), while the current data only represent a single geological stage.

### Neural Network configuration

We custom-designed a BNN classification model that maps raw geographic and temporal distances of selected taxon occurrences (fossil or current) to a set of vegetation classes. These distance features can be complemented by any set of additional features, such as the abiotic features used in this study. The BNN model consists of multiple hidden layers generating a numerical representation of the features in multidimensional space, as well as an output layer that maps the nodes of the last hidden layer to the output classes, in this case open and closed habitats. Given the flexibility of our model and the fact that it is based on absolute distance measures, it may be applied to any vegetation prediction task, independently of the spatial and temporal scale of the project.

The first two hidden layers are only applied to the taxon distance features rather than to the additional abiotic features. In these layers the raw spatial and temporal occurrence distances are combined into a single value per taxon, which represents a measure of proximity of each taxon to a given input instance. The raw distances are provided in pairs of one spatial and one temporal distance measurement, both associated with a specific occurrence of a taxon (Fig. 1A). In the first layer these two distances are merged into one spatiotemporal distance value via matrix multiplication with a space and a time weight (Fig. 1B). To reduce the number of estimated parameters for better convergence, the space and time weights are shared among all occurrences under the assumption that the relative importance of space and time in determining the proximity of a given occurrence is expected to be the same for all occurrences of different taxa. After collapsing spatial and temporal distances into one spatiotemporal distance value in this way, we estimate specific taxon-weights for each taxon and geological stage, which are then used to collapse the multiple spatiotemporal distances across different geological stages into one single feature value for each taxon.

Depending on the chosen pooling strategy, these taxon feature values are either fed as individual features into the next layer (no pooling) or are summarized into one faunal and one floral feature, by either extracting the maximum output value (max-pooling) or by summing all output values (sum-pooling) across all mammal and plant taxa, respectively. Following the initial two layers, the taxon-features (n=100 or n=2, depending on pooling strategy) are fed together with the additional abiotic features (n=6) into a fully connected neural network and eventually mapped to the binary vegetation classes in the output layer (Fig. 1C). We tested different network configurations in terms of number of layers and nodes per layer, different pooling strategies, as well as different combinations of training features and instances, and selected the best model based on the highest test set prediction accuracy (Table 1).

During training, all weights of the model are initially drawn randomly from a normal distribution centered in 0 and are then updated and sampled using a Metropolis Hastings Markov Chain Monte Carlo (MCMC) algorithm ^41^. We used a standard normal prior for all weights (parameters of the model). Further, we applied a ReLU activation function for each hidden layer, and a softmax function for the output layer. The likelihood is calculated as the cumulative loss across all training instances. For all models we ran an MCMC chain for 1 million generations, sampling every 200 iterations.

Our BNN implementation allows not only to estimate the most probable vegetation label for a given point in time and space, but also to calculate the posterior probability of this label, providing an inherent measure of uncertainty. We calculated the posterior probability of each class label for a given instance as the mean class probability across all posterior samples. This ability makes BNNs an attractive alternative to regular neural network algorithms, which allow no such uncertainty modeling, although analogous approximations exist, such as Monte Carlo dropout ^42^.

#### Feature importance

To determine the relative importance of each feature used in our model, we applied the method of permutation feature importance (sensu Breiman ^43^). In this approach, the values of a given feature are randomly shuffled across all instances of the test set. This process masks any existing information that lies within the values of a given feature. The class labels for all test instances are then predicted using the modified feature matrix. The resulting test accuracy is then compared with that of the original feature matrix and the difference between these accuracies (*Δacc*) is interpreted as a measure of relative importance of the shuffled feature for the classification task. We repeated this process for each feature column in our feature matrix (n=106) and ranked the features based on their *Δacc* values (Fig. 4).

### Predicting vegetation labels

To produce continuous vegetation maps across North America, we constructed a 0.5° x 0.5° grid across the cropping window defined in this study and extracted the coordinates of the cell-center for each grid cell. For these points, we extracted spatiotemporal taxon-distances and abiotic features in the same manner as for the training instances. We repeated this process in 1 Ma steps starting in the present (t=0) throughout the last 30 Ma (t=30), producing 31 feature-datasets of North America through time, considering tectonic movement (*mapast* ^24^). Based on the BNN weights sampled during training by the MCMC (excl. 10% burn-in) we determined the posterior probabilities of each vegetation label for each given point (Fig. 2).

## Acknowledgements

We thank all paleobotanists for producing and making publicly available the paleovegetation data necessary for training the models presented in this study. Further we thank Juan Carrillo for providing paleontological feedback, in particular on the mammal fossil data. TA and DS received funding from the Swedish Research Council (VR: 2019-04739). DS received funding from the Swiss National Science Foundation (PCEFP3_187012; FN-1749).

## Supplementary Material

The supplementary material accompanying this study contains the following:

- Supplementary Results
- Supplementary Table S1
- Supplementary Figures S1-S5

Additionally, Supplementary Data (S1 and S2) can be downloaded at https://tinyurl.com/pd6f2n3k.

## References

1. Lu, Z. et al. Vegetation Pattern and Terrestrial Carbon Variation in Past Warm and Cold Climates. Geophysical Research Letters 46, 8133–8143 (2019).

2. Peppe, D. J. Megafloral change in the early and middle Paleocene in the Williston Basin, North Dakota, USA. Palaeogeography, Palaeoclimatology, Palaeoecology 298, 224–234 (2010).

3. Janis, C. M. A climatic explanation for patterns of evolutionary diversity in ungulate mammals. Palaeontology 32, 463–481 (1989).

4. Carvalho, M. R. et al. Extinction at the end-Cretaceous and the origin of modern Neotropical rainforests. Science 372, 63–68 (2021).

5. Niklas, K. J., Tiffney, B. H. & Knoll, A. H. Patterns in vascular land plant diversification. Nature 303, 614–616 (1983).

6. Condamine, F. L., Silvestro, D., Koppelhus, E. B. & Antonelli, A. The rise of angiosperms pushed conifers to decline during global cooling. PNAS 117, 28867–28875 (2020).

7. Silvestro, D. et al. Fossil data support a pre-Cretaceous origin of flowering plants. Nat Ecol Evol 5, 449–457 (2021).

8. Edwards, E. J., Osborne, C. P., Strömberg, C. A. E., Smith, S. A. & Consortium, C. G. The Origins of C4 Grasslands: Integrating Evolutionary and Ecosystem Science. Science 328, 587–591 (2010).

9. Strömberg, C. A. Evolution of grasses and grassland ecosystems. Annual review of Earth and planetary sciences 39, 517–544 (2011).

10. Gibson, D. J. Grasses and Grassland Ecology. (Oxford University Press, 2009).

11. Miller, L., Smith, S., Sheldon, N. & Stromberg, C. Eocene vegetation and ecosystem fluctuations inferred from a high-resolution phytolith record. Geological Society of America Bulletin 124, 1577–1589 (2012).

12. Fox, D. L. et al. Climatic Controls on C4 Grassland Distributions During the Neogene: A Model-Data Comparison. Front. Ecol. Evol. 6, (2018).

13. Strömberg, C. A. E., Dunn, R. E., Crifò, C. & Harris, E. B. Phytoliths in Paleoecology: Analytical Considerations, Current Use, and Future Directions. in Methods in Paleoecology: Reconstructing Cenozoic Terrestrial Environments and EcologicalCommunities (eds. Croft, D. A., Su, D. F. & Simpson, S. W.) 235–287 (Springer International Publishing, 2018). doi:10.1007/978-3-319-94265-0_12.

14. Jaramillo, C. 140 million years of tropical biome evolution. The Geology of Colombia (ed. Gomez, J.). Colombian Geological Survey, Bogota, Colombia (2019).

15. Barbolini, N. et al. Cenozoic evolution of the steppe-desert biome in Central Asia. Science Advances 6, eabb8227 (2020).

16. Jaramillo, C. & Cárdenas, A. Global warming and neotropical rainforests: A historical perspective. Annual Review of Earth and Planetary Sciences 41, 741–766 (2013).

17. Kaplan, J. O. Geophysical Applications of Vegetation Modeling. Infoscience https://infoscience.epfl.ch/record/136645 (2001).

18. Pound, M. J. et al. A Tortonian (Late Miocene, 11.61–7.25Ma) global vegetation reconstruction. Palaeogeography, Palaeoclimatology, Palaeoecology 300, 29–45 (2011).

19. MacFadden, B. J. Origin and evolution of the grazing guild in new world terrestrial mammals. Trends in Ecology & Evolution 12, 182–187 (1997).

20. Jacobs, B., Kingston, J. & Jacobs, L. The Origin of Grass-Dominated Ecosystems. Annals of the Missouri Botanical Garden 86, 590 (2000).

21. Dunn, R. E., Strömberg, C. A. E., Madden, R. H., Kohn, M. J. & Carlini, A. A. Linked canopy, climate, and faunal change in the Cenozoic of Patagonia. Science 347, 258–261 (2015).

22. Alroy, J., Marshall, C. & Miller, A. Paleobiology database. (NCEAS, 2004).

23. Scotese, C. R., Song, H., Mills, B. J. & van der Meer, D. G. Phanerozoic paleotemperatures: The earth’s changing climate during the last 540 million years. Earth-Science Reviews 103503 (2021).

24. Varela, S. & Rothkugel, K. S. mapast: combine paleogeography and paleobiodiversity. (2018).

25. Zachos, J. C., Dickens, G. R. & Zeebe, R. E. An early Cenozoic perspective on greenhouse warming and carbon-cycle dynamics. nature 451, 279–283 (2008).

26. Jung, M., Henkel, K., Herold, M. & Churkina, G. Exploiting synergies of global land cover products for carbon cycle modeling. Remote Sensing of Environment 101, 534–553 (2006).

27. Strömberg, C. A. & McInerney, F. A. The Neogene transition from C3 to C4 grasslands in North America: assemblage analysis of fossil phytoliths. Paleobiology 37, 50–71 (2011).

28. Leopold, E. B., GengWu, L. & Clay-Poole, C. Low-biomass vegetation in the Oligocene? in Eocene-Oligocene climatic and biotic evolution 399–420 (Princeton Univ. Press, 1992).

29. Hembree, D. I. & Hasiotis, S. T. Paleosols and ichnofossils of the White River Formation of Colorado: Insight into soil ecosystems of the North American Midcontinent during the Eocene-Oligocene transition. Palaios 22, 123–142 (2007).

30. Ehleringer, J. R., Cerling, T. E. & Helliker, B. R. C4 photosynthesis, atmospheric CO2, and climate. Oecologia 112, 285–299 (1997).

31. Sandom, C., Ejrnæs, R., Hansen, M. & Svenning, J.-C. High herbivore density associated with vegetation diversity in interglacial ecosystems. Proceedings of the National Academy of Sciences of the United States of America 111, (2014).

32. Jeffers, E. S. et al. Plant controls on Late Quaternary whole ecosystem structure and function. Ecology Letters 21, 814–825 (2018).

33. Cohen, K. M., Finney, S. C., Gibbard, P. L. & Fan, J.-X. The ICS international chronostratigraphic chart. Episodes 36, 199–204 (2013).

34. Fortelius, M. New and Old Worlds database of fossil mammals (NOW). (University of Helsinki, Helsinki, Finland, 2013).

35. Williams, J. W. et al. The Neotoma Paleoecology Database, a multiproxy, international, community-curated data resource. Quaternary Research 89, 156–177 (2018).

36. Carrasco, M. A., Barnosky, A. D., Kraatz, B. P. & Davis, E. B. The Miocene Mammal Mapping Project (Miomap): An Online Database of Arikareean Through Hemphillian Fossil Mammals. Bulletin of Carnegie Museum of Natural History 2007, 183–188 (2007).

37. Silvestro, D., Antonelli, A., Salamin, N. & Meyer, X. Improved estimation of macroevolutionary rates from fossil data using a Bayesian framework. bioRxiv 316992 (2018) doi:10.1101/316992.

38. Xing, Y. et al. Testing the biases in the rich Cenozoic angiosperm macrofossil record. International Journal of Plant Sciences 177, 371–388 (2016).

39. GBIF. GBIF Home Page. GBIF Home Page https://www.gbif.org (2020).

40. Chamberlain, S. A. & Boettiger, C. R Python, and Ruby clients for GBIF species occurrence data. https://peerj.com/preprints/3304 (2017) doi:10.7287/peerj.preprints.3304v1.

41. Silvestro, D. & Andermann, T. Prior choice affects ability of Bayesian neural networks to identify unknowns. arXiv preprint 2005.04987, (2020).

42. Gal, Y. & Ghahramani, Z. Dropout as a bayesian approximation: Representing model uncertainty in deep learning. in international conference on machine learning 1050–1059 (2016).

43. Breiman, L. Random forests. Machine learning 45, 5–32 (2001).

