## Supplementary Material for "The evolution of open habitats in North America revealed by deep learning models"

This supplementary material file contains the following:

- Supplementary Results
- Supplementary Table S1
- Supplementary Figures S1-S5

Additionally, Supplementary Data (S1 and S2) can be downloaded at https://tinyurl.com/pd6f2n3k.

### Supplementary Results

#### Evaluating prediction accuracy on current vegetation map

For the task of predicting paleovegetation, which is presented in the main text, we only have limited data points with past vegetation information (those compiled through the literature review). This limits our ability to evaluate how accurately the trained models predict the true vegetation patterns, based on these few subsampled data points. Yet, the current vegetation data for North America provides a large dataset and thus a suitable framework to test the ability to predict the overall vegetation pattern correctly.

To investigate how well our model can predict vegetation only based on a small subsample of the actual vegetation, we trained a BNN model using only 281 of the approximately 11,000 current vegetation points across North America (Supplementary Fig. S3 A+B). This number of training points is equal to the number of paleovegetation points used for training of the paleo-models, although in case of the paleo-data these points are distributed throughout several geological stages. While these training data only constitute about 2.5% of the current vegetation information, our model was able to predict the entire current vegetation (Supplementary Fig. S3 C), with a prediction accuracy of 88.1% (Table 1). When making visible where on the map our model misclassified vegetation labels, we find that these misclassifications predominantly occur in the transitioning areas between open and closed vegetation (Supplementary Fig. S3 D). This suggests that, while our model accurately learns the general distributions of both vegetation types, the misclassifications are mainly a result of limited resolution of the vegetation boundaries, likely caused by the relatively small number of training vegetation points.

We find that a 5-fold increase in the number of training points (n=1,405) increases the resolution of the predictions, leading to an increased prediction accuracy of 91.6% (Table 1, Supplementary Fig. S4). This shows that more data points can increase the accuracy of the model, suggesting that our model accuracies presented in the main text could be further improved with more paleovegetation data points being available. However, this bottleneck of limited training data and the resulting limitations in prediction accuracy can be partly accommodated by applying posterior thresholds to the vegetation predictions (see below).

#### Applying posterior thresholds to increase accuracy

One of the advantages of using BNNs over regular neural network implementations is the explicit modeling of uncertainty in the model predictions by producing posterior probability (PP) estimates for each vegetation prediction (see Methods for more detailed information). These PPs are derived from a posterior sample of the BNN weights and are thus different in their nature to the class-probabilities resulting from the output layer of a regular neural network, which instead represent point estimates. We utilize the PP estimates by setting a posterior threshold to only make vegetation predictions for sites that have a high prediction certainty, while assigning all predictions below the thresholds as “unknown”. We select this PP threshold individually for each model to ensure a specified minimum prediction accuracy.

In the case of our model trained on 281 current vegetation points, we find that a PP threshold of 0.56 leads to an expected prediction accuracy of > 90% (Table 1), while still allowing to make vegetation predictions for 94.5% of all North American terrestrial cells (Supplementary Fig. S5). Increasing the PP threshold to 0.74 leads to an expected prediction accuracy of > 95%, while predicting 77.2% of the map with high confidence (Supplementary Fig. S3 E). For the model trained on 1,405 current vegetation labels, a posterior threshold of 0.66 was sufficient to ensure 95% prediction accuracy, allowing to make high accuracy vegetation predictions for 89.5% of North American terrestrial cells (Supplementary Fig. S4). Prediction accuracies even higher than 95% could be achieved by setting higher posterior thresholds, at the cost of an increasing number of predictions labeled as unknown (Supplementary Fig. S5). This trade-off between quality and quantity of the predictions allows us to focus the predictions of the model only on areas for which we can confidently infer vegetation, leading to a substantial increase in prediction accuracy (Supplementary Fig. S3 F). This property of BNN models is of particular interest when the number of available vegetation information for training is limited, as is the case for our paleovegetation models.

### Supplementary Tables

**Supplementary Table S1**. Association scores of mammal and plant taxa with open habitat. The openness score was calculated as the fraction of occurrences of each taxon that fall within predicted open habitat, averaged across the predicted 1 Mya increments throughout the last 30 Mya.

| Taxon | Openness score | Taxon | Openness score | Taxon | Openness score |
|---|---|---|---|---|---|
| Cynomys | 0.99 | Puma | 0.55 | Lauraceae | 0.27 |
| Onychomys | 0.86 | Tapirus | 0.55 | Platanaceae | 0.24 |
| Ochotona | 0.80 | Equus | 0.55 | Gomphotherium | 0.23 |
| Perognathus | 0.79 | Ondatra | 0.55 | Erethizon | 0.22 |
| Zapus | 0.78 | Castor | 0.51 | Sylvilagus | 0.21 |
| Brachyerix | 0.75 | Taxidea | 0.51 | Fabaceae | 0.21 |
| Baiomys | 0.74 | Cryptotis | 0.49 | Myricaceae | 0.20 |
| Neotoma | 0.74 | Mammuthus | 0.49 | Eumops | 0.20 |
| Mustela | 0.73 | Phenacomys | 0.48 | Panthera | 0.20 |
| Mephitis | 0.69 | Anacardiaceae | 0.47 | Lontra | 0.19 |
| Scalopus | 0.68 | Lepus | 0.46 | Typhaceae | 0.17 |
| Tamias | 0.68 | Lasionycteris | 0.46 | Ericaceae | 0.17 |
| Aphelops | 0.67 | Boraginaceae | 0.45 | Oleaceae | 0.15 |
| Tayassu | 0.67 | Peromyscus | 0.45 | Juglandaceae | 0.15 |
| Notiosorex | 0.66 | Meliaceae | 0.45 | Ulmaceae | 0.15 |
| Reithrodontomys | 0.66 | Thomomys | 0.44 | Berberidaceae | 0.15 |
| Oryzomys | 0.66 | Dipodomys | 0.44 | Asteraceae | 0.15 |
| Sorex | 0.65 | Mammut | 0.43 | Rosaceae | 0.14 |
| Spermophilus | 0.65 | Urocyon | 0.43 | Polygonaceae | 0.14 |
| Dipoides | 0.63 | Ursus | 0.42 | Vitaceae | 0.14 |
| Blarina | 0.63 | Geomys | 0.42 | Salicaceae | 0.13 |
| Hypolagus | 0.62 | Ranunculaceae | 0.41 | Cyperaceae | 0.13 |
| Ammospermophilus | 0.61 | Eptesicus | 0.39 | Grossulariaceae | 0.12 |
| Poaceae | 0.61 | Marmota | 0.38 | Sapindaceae | 0.12 |
| Myotis | 0.61 | Procyon | 0.38 | Fagaceae | 0.11 |
| Vulpes | 0.61 | Ebenaceae | 0.37 | Dasypus | 0.10 |
| Cannabaceae | 0.60 | Sigmodon | 0.35 | Betulaceae | 0.09 |
| Bassariscus | 0.60 | Platygonus | 0.35 | Caprifoliaceae | 0.06 |
| Canis | 0.59 | Odocoileus | 0.34 | Cornaceae | 0.03 |
| Antrozous | 0.59 | Sciurus | 0.32 | Potamogetonaceae | 0.02 |
| Spilogale | 0.57 | Felis | 0.30 | Nymphaeaceae | 0.02 |
| Cratogeomys | 0.56 | Adoxaceae | 0.30 | Altingiaceae | 0.00 |
| Scapanus | 0.56 | Hydrangeaceae | 0.29 | | |
| Lynx | 0.55 | Rhamnaceae | 0.29 | | |

### Supplementary Figures

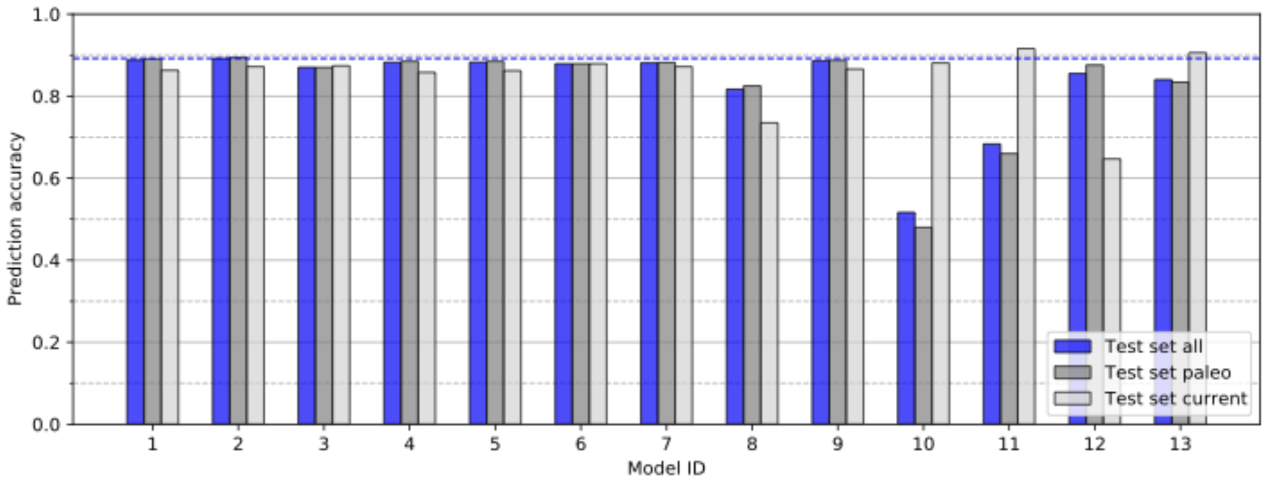


**Supplementary Figure S1.** Prediction accuracy of tested models. The tested scenarios differed in the number of nodes (32-8 or 8), the used features (all, only biotic, or only abiotic), the pooling strategy applied to the biological features (sum-pooling or max-pooling), and the number of current vegetation instances used for training (0, 281, or 1405, see Table 1 in main text for model ID key). Blue bars show the overall prediction accuracy of each model (five-fold cross validation), which constitutes the weighted mean of the paleovegetation prediction accuracy (dark grey) and the prediction accuracy of for the entire North American current vegetation map (light grey), excluding the grid cells that were used for training. The blue horizontal bar shows the maximum reached prediction accuracy of 89.2% (model #2).

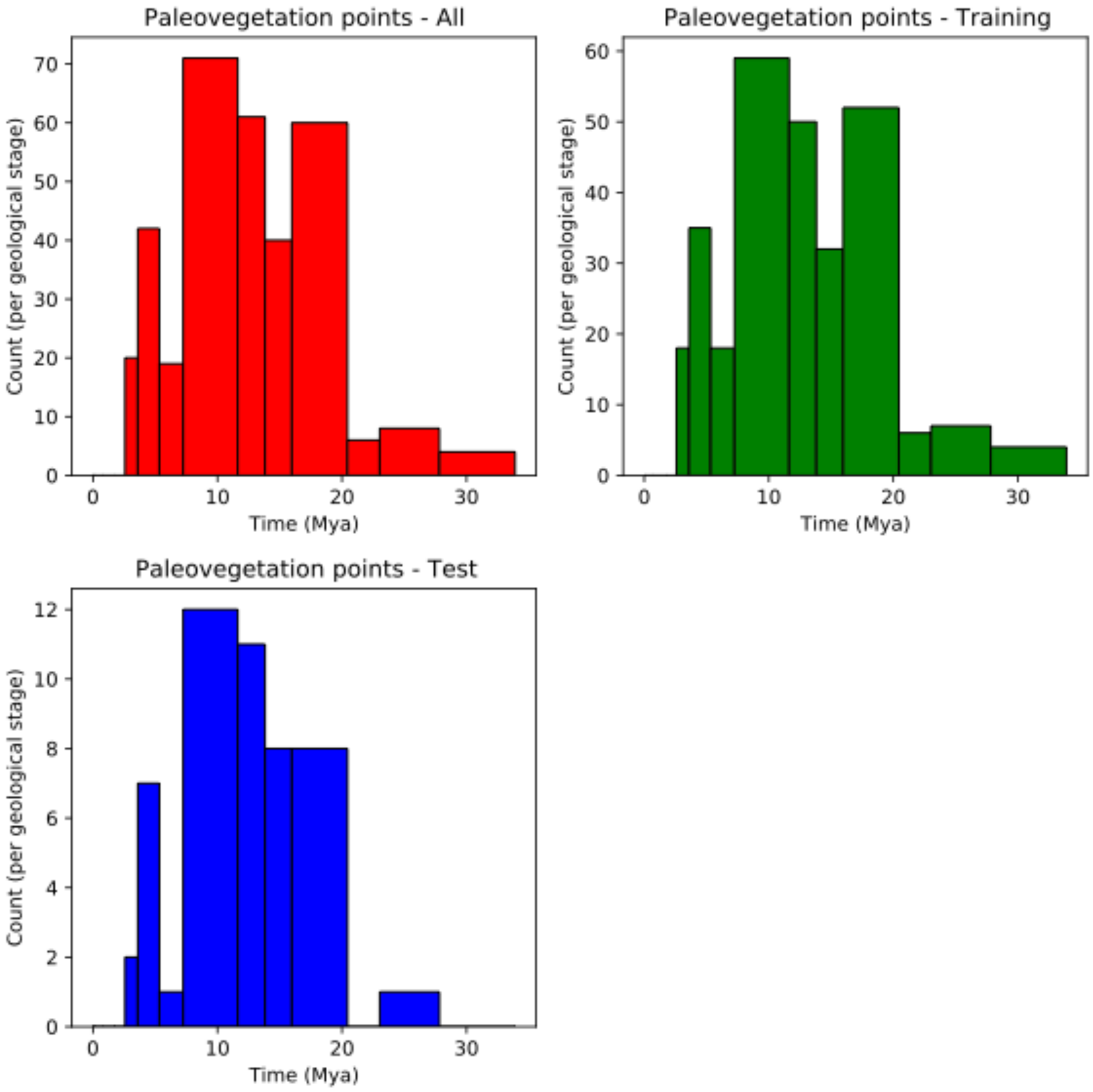


**Supplementary Figure S2.** Temporal distribution of paleovegetation sites analyzed. The ages shown in the histograms were binned using the geological stage boundaries according to the International Chronostratigraphic Chart, v2020/03.

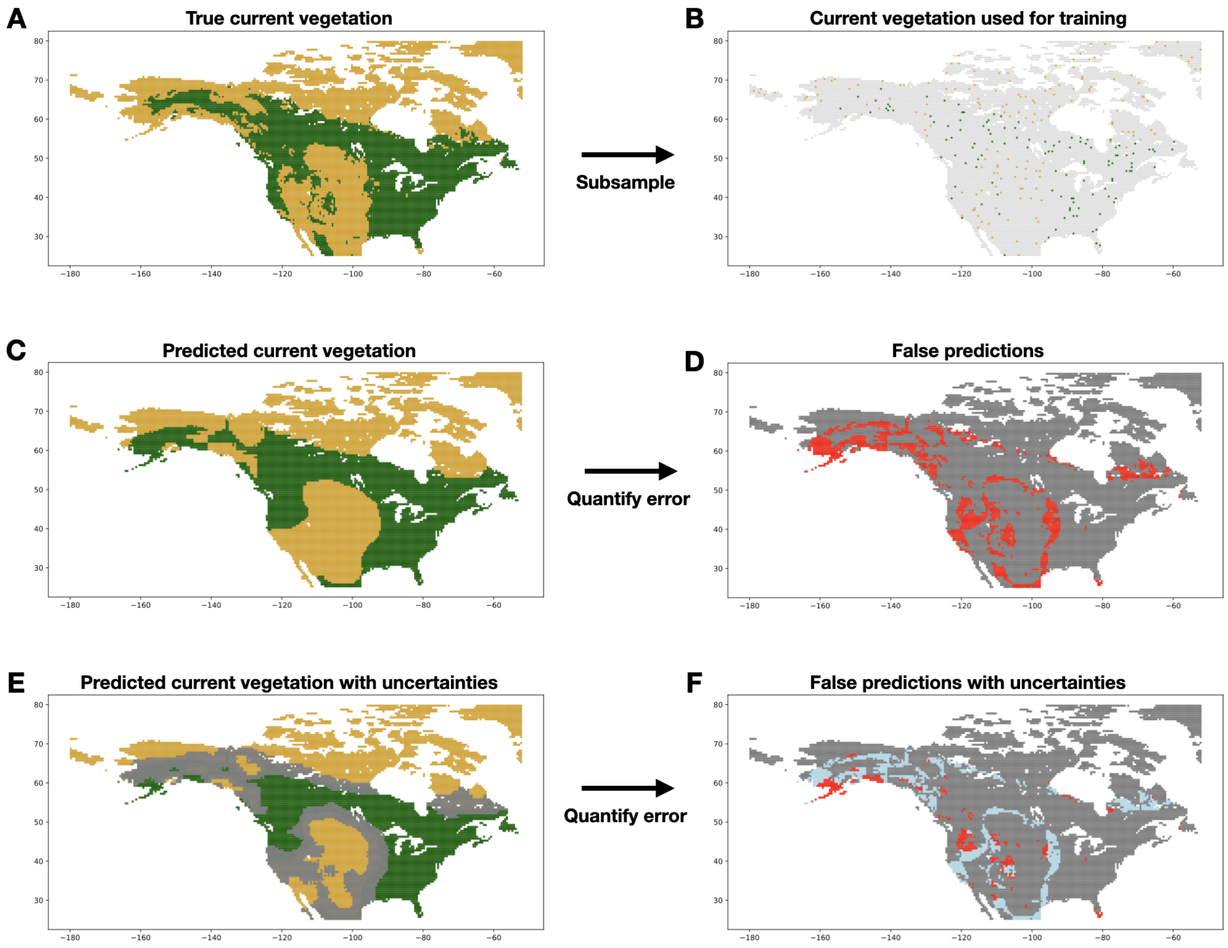


**Supplementary Figure S3.** Utility of applying posterior probability thresholds to vegetation predictions. The true current distribution of open (yellow) and closed habitat (green) in North America (A) is based on the SYNMAP potential vegetation data. A random subsample of 281 of these current vegetation points (B) was used for training the BNN model. We used the trained model to predict the current vegetation of North America, based on the mammal, plant, and climate associations with the vegetation type that were learned by the model (C). While the majority of the current map was predicted correctly (84.8% accuracy, excl. training points), there are several incorrect vegetation predictions, highlighted in red (D). A posterior threshold can be applied to our model predictions (E), which allows us to distinguish between confident vegetation predictions (colored), and those the model is uncertain about (grey). In this case we applied a posterior threshold that ensures a prediction accuracy of > 95%. The resulting predictions show only a small fraction of falsely predicted vegetation labels (F), while the majority of the problematic predictions are now modeled as uncertain (light blue).

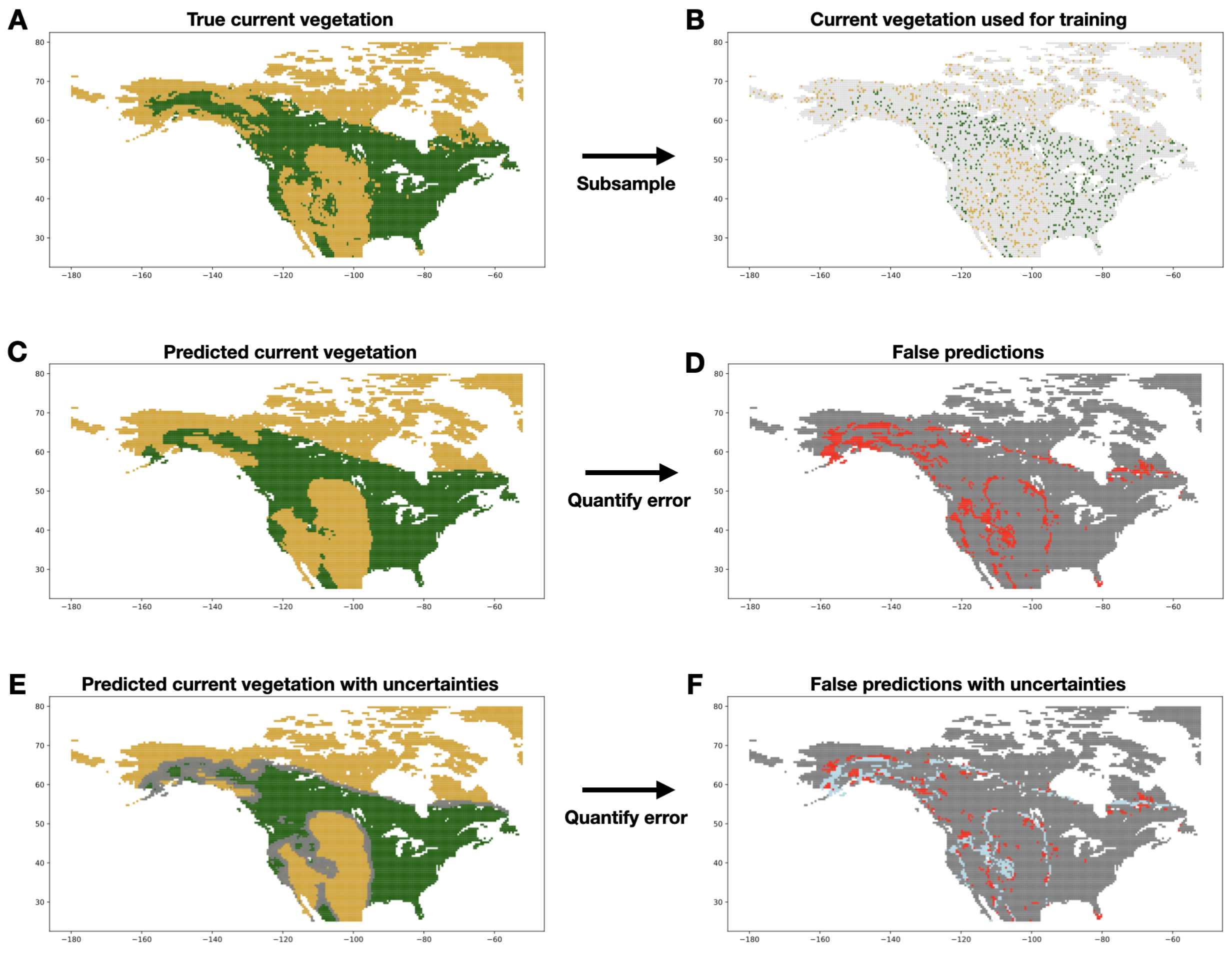


**Supplementary Figure S4**. Predictions of a model trained with 1,405 current vegetation sites. See caption of Supplementary Fig. S3 for more details.

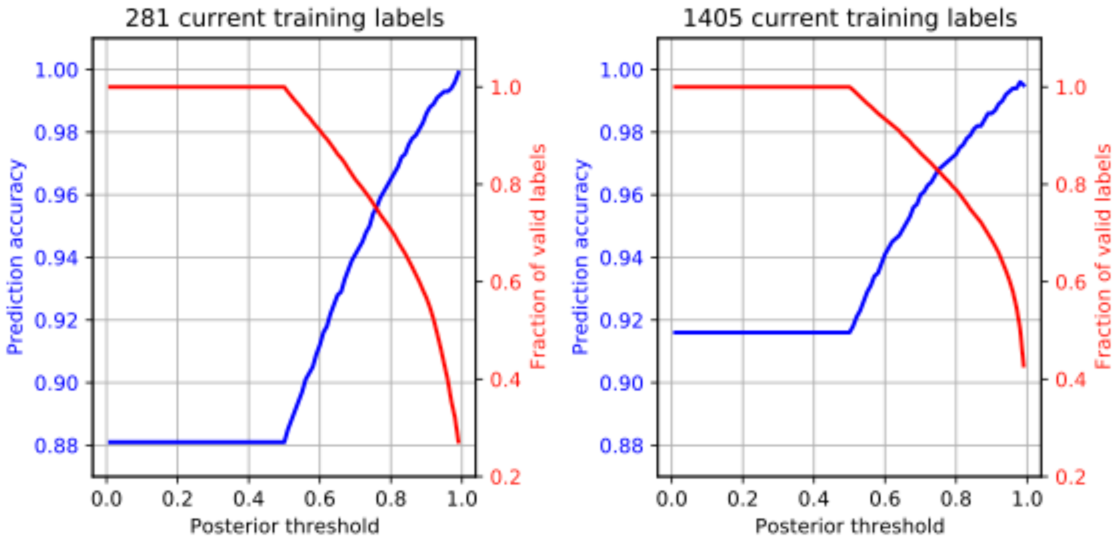


**Supplementary Figure S5**. Trade-off between increasing prediction accuracy and fewer vegetation label predictions with increasingly strict posterior thresholds. Results are shown for a model trained on 281 current vegetation labels (no paleontological data), as well as for a model with a 5-fold increased number of current training labels (n=1,405). The displayed accuracies were calculated based on all current vegetation labels across North America, excluding the labels used for training.
